# Diaphragmatic Central Motor Conduction Changes In Chronic Obstructive Pulmonary Disease

**DOI:** 10.1101/646836

**Authors:** Rehab Abdelaal El-Nemr, Rania Ahmad Sweed, Hanaa Shafiek

## Abstract

**Background and objectives:** Respiratory muscles dysfunction has been reported in COPD. Transcranial magnetic stimulation (TMS) is easy non-invasive that has been used for assessing the respiratory corticospinal pathways particularly of diaphragm. We aimed to study the cortico-diaphragmatic motor system changes in COPD using TMS and to correlate the findings with the pulmonary function.

**Methods:** A case control study recruited 30 stable COPD from the out-patient respiratory clinic of Main Alexandria University hospital-Egypt and 17 healthy control subjects who were subjected to spirometry. Cortical conduction of the diaphragm was performed by TMS to all participants followed by cervical magnetic stimulation of the phrenic nerve roots. Diaphragmatic resting motor threshold (DRMT), cortical motor evoked potential latency (CMEPL), CMEP amplitude (CMEPA), peripheral motor evoked potential latency (PMEPL), PMEP amplitude (PMEPA) and central motor conduction time (CMCT) were measured.

**Results:** 66.7% of COPD patients had severe and very severe COPD with median age of 59 (55-63) years. There was statistically significant bilateral decrease in DRMT, CMEPA and PMEPA in COPD group versus healthy subjects and significant increase in CMEPL and PMEPL (*p* <0.01). Left CMCT was significantly prolonged in COPD group versus healthy subjects (*p* <0.0001) but not right CMCT. Further, there was significant increase in CMEPL and CMCT of left versus right diaphragm in COPD group (*p*= 0.003 and 0.001 respectively) that inversely correlated with FEV_1_% and FVC% predicted.

**Conclusion:** Central cortico-diaphragmatic motor system is affected in COPD patients with heterogeneity of both sides that is correlated with pulmonary function.

**Significance:** Coticospinal pathway affection could be a factor for development of diaphragmatic dysfunction in COPD patients accordingly its evaluation could help in personalization of COPD management especially pulmonary rehabilitation programs

## Introduction

Chronic obstructive pulmonary disease (COPD) is mainly presented with dyspnea and exercise limitation secondary to irreversible airflow obstruction; however, nowadays COPD is considered as multi-systemic inflammatory disorder rather than simple respiratory disease.[1] Respiratory muscles dysfunction has been reported in COPD compared to healthy elderly individuals [2] and has been implicated in the development of dyspnea.

Respiratory muscles, particularly the diaphragm which is considered the main inspiratory muscle, are affected in COPD in two main ways. Firstly, change of shape and geometry of the chest wall secondary to air trapping and hyperinflation in COPD leads to chronic reduction of the apposition zone of the diaphragm [3] and shorten of the diaphragm fiber sarcomere. [4] Secondly, local activation of muscle proteases and oxidative stress due to inspiratory loading induce structural muscular injury [5, 6] that is further affected by exposure to systemic inflammatory process associated with COPD. [7]

Transcranial magnetic stimulation (TMS) is easy, non-invasive and painless tool that aimed at measuring neuronal electrical activity. [8] TMS has been used as investigation tool for assessing the respiratory corticospinal pathways and studying of diaphragmatic motor evoked potential (MEP). [8–12] TMS has been used to identify central origin of a diaphragmatic dysfunction in stroke, [13] multiple sclerosis, [14] amyotrophic lateral sclerosis, [15] or spinal cord injury. [16]

Cervical magnetic stimulation of the phrenic nerve roots has been used previously to assess diaphragm weakness in COPD patients. [17] In the last decade, a recent study demonstrated increased excitability of the motor cortex controlling respiratory muscles in COPD especially diaphragm which could be secondary to increased inspiratory load and subsequent elevated respiratory drive. [18] However, still little research has been conducted in COPD to assess central neural drive to the diaphragm and its possible involvement in physiological derangement in COPD patients. Accordingly, we aimed to study the cortico-diaphragmatic motor system changes in COPD patients using TMS; to correlate the findings with the pulmonary function; and to detect possible cutoff value for corticospinal diaphragmatic pathway affection that could be a reference in this group of patients.

## Methods

### Study design and ethics

A case control study recruited 30 stable COPD according to updated GOLD guidelines 2017 [1] who attended the out-patient respiratory clinic of Main Alexandria University hospital, Egypt as well as 17 healthy control subjects who were invited to participate in the study. The study has been approved by the scientific committee of faculty of medicine, Alexandria University, Egypt. A written informed consent was obtained from all participants enrolled in this study. The study was conducted over 10 months.

### Study population and their characterization

Thirty COPD patients were included in the study. All patients were stable i.e. no COPD exacerbation in last 4 weeks, and has been proved to have airway obstruction using spirometry (post-bronchodilator FEV_1_/FVC < 0.70) as being accepted by updated GOLD guidelines 2017. [1] All patients who were known to have COPD exacerbation, current oral corticosteroids therapy or within last 30 days, bronchial asthma, interstitial lung diseases, metabolic diseases (mainly diabetes mellitus, uremia and hepatic failure), neurological diseases (as cerebrovascular stroke, epilepsy, peripheral neuropathy and muscle diseases), body mass index (BMI) more than 40 kg/mm^2^, history of drug abuse, history of any neoplasm, or any contraindications for magnetic stimulation were excluded from the study. Further, 17 healthy control subjects with normal lung function referred for check-up were recruited from other clinics.

All the participants underwent detailed history taking that included age, sex, smoking history, respiratory symptoms, current medications; followed by local and general examination, chest X-ray, spirometry for measurement of post-bronchodilator FVC, FEV_1_ and FEV_1_/FVC ratio. For COPD patients, arterial blood gases (ABG) were assessed for COPD patients, and venous blood sample was taken for measurement of fasting blood glucose, liver function testing, renal function testing, complete blood picture, and serum electrolytes (sodium and potassium). Computed tomography of chest was performed if indicated clinically.

### Diaphragmatic neural function assessment

Firstly, TMS of the diaphragm was carried out using electrophysiological apparatus with a circular coil (Nihon Kohden MEB-7102K© with peak magnetic field strength of 2 Tesla; Tokyo, Japan). The coil was applied tangentially to the scalp of patient at diaphragmatic motor cortical area, a point of optimal excitability, located 3 cm lateral to midline and 2-3 cm anterior to auricular plane [9] with face A of the coil visible from above for left hemisphere stimulation and face B for right hemisphere stimulation recording cortical MEPs responses. Surface electrodes were placed in the 7^th^ and 8^th^ right and left intercostal spaces respectively within the anterior axillary line, and the reference electrode on the corresponding lower rib for recording diaphragmatic cortical MEP response contralateral to the stimulation site. A ground electrode was placed on the manubrium sterni. [19] The recording conditions utilized were: filter setting high at 3K Hz and low at 3Hz, vertical gain 0.2-2mV/ division, and sweep speed 5 msec/division. The stimulus threshold was determined by increasing the stimulus strength (expressed as the % of the maximum output of the stimulator) until 3 reproducible MEPs responses were obtained, then the stimulus intensity was set at 20% above the threshold value. The angle of the coil around the stimulation site was changed until the highest MEP response during inspiratory phase was recorded. The following parameters were measured from central stimulation: diaphragmatic resting motor threshold (DRMT), cortical motor evoked potential latency (CMEPL) in milliseconds (ms), CMEP amplitude (CMEPA) in microvoltage (μv).

Secondly, cervical magnetic stimulation of the phrenic nerve roots in the neck was performed bilaterally. The periphery of the circular coil of the apparatus was placed 2 cm lateral to mid-line and 1-2 cm above the 5^th^ cervical spine while the patient head slightly bent forward. The diaphragmatic peripheral motor evoked potential (PMEP) was recorded by using the same recording electrodes setting previously discussed whereas peripheral motor evoked potential latency (PMEPL) and PMEP amplitude (PMEPA) were measured. Central motor conduction time (CMCT) was then calculated as follow: CMCT = CMEPL – PMEPL. [20]

### Statistical analysis

Quantitative data were expressed as mean ± standard deviation (SD) or median (interquartile range; 25-75 percentile) according to the normal distribution of data while qualitative data was expressed as number and percentage (%). Mann - Whitney test, Kruskal - Wallis test, student t-test, Chi-square test and Spearman rho correlation were used as appropriate. ROC (receiver operating characteristic) curve and area under the curve (AUC) has been used to detect cutoff values for diaphragmatic CMEPs that could differentiate COPD from healthy individuals. All the analysis has been performed using MedCalc^®^ (version 9.2.1.0, Acacialaan 22, B-8400 Ostend, Belgium) and SPSS package (PASW Statistics for Windows, Version 22.0. Chicago: SPSS Inc.).

## Results

### Participants’ characteristics

All the baselines characteristics of COPD patients and healthy control are shown in table “1”. All the recruited patients were males with no statistically significant difference between both groups regarding age, BMI, and smoking status; however smoking index was significantly higher in COPD group (*p* < 0.0001). Baseline FVC%, FEV_1_% and FEV_1_/FVC were significantly lower in COPD group (*p* < 0.0001) whereas 2 COPD patient (6.7%) had mild airway obstruction, 8 patients (26.7%) had moderate airway obstruction, 12 patients (40%) had severe airway obstruction and 8 patients (26.7%) had very severe airway obstruction according GOLD classification.

**Table (1):** Demographic and baseline clinical characteristics of study population

| **Character** | **COPD (n=30)** | **Control (n=17)** | ***p* value** |
| --- | --- | --- | --- |
| Age (years) | 59 (55- 63) | 55 (50 - 59.5) | 0.055 |
| **Gender**<br>Male / Female | 30 (100) / 0 (0) | 17 (100) / 0 (0) | 1.0 |
| BMI (Kg/mm<sup>2</sup>) | 24.3 ± 4.7 | 22.8 ± 3.6 | 0.338 |
| **Smoking history**<br>smoker / ex-smoker<br>smoking index (PYI) | 14 (46.7) / 16 (53.3)<br>60 (45 - 80) | 11 (64.7) / 6 (35.3)<br>20 (10-30) | 0.375<br><0.0001* |
| **Comorbidities**<br>Hypertension<br>IHD<br>Obesity and OSA | 10 (33.3)<br>6 (20)<br>1 (3)<br>3 (10) | 0 (0) | 0.029* |
| **Spirometry**<br>FVC% predicted | 56 (50.3 - 66.3) | 109 (98 - 123) | < 0.0001* |
| FEV<sub>1</sub>% predicted | 42.9 (29 - 54) | 123 (112 - 136.5) | < 0.0001* |
| FEV<sub>1</sub>/FVC | 57.6 ± 8.7 | 86.6 ± 8.5 | < 0.0001* |
| **ABG**<br>pH<br>PaO<sub>2</sub> (mmHg)<br>PaCO<sub>2</sub> (mmHg)<br>HCO<sub>3</sub> (mmol/L)<br>SaO<sub>2</sub> | 7.43 ± 0.048<br>78.43 ± 20.8<br>40.5 ± 8.9<br>25 (22 - 30)<br>96 (94.8 - 97.0) | NA | NA |
| **Laboratory tests** |  | NA | NA |
| FBS (mg/dl) | 101.5 (72 - 111) |  |  |
| Hb (g/dl) | 13.9 ± 1.3 |  |  |
| BUN (mg/dl) | 15 (12 - 20) |  |  |
| Cr (mg/dl) | 0.81 ± 0.24 |  |  |
| Na (mmol/L) | 140 (137 - 144) |  |  |
| K (mmol/L) | 4.1 ± 0.35 |  |  |
| AST (U/L) | 29.5 (22 - 41) |  |  |
| ALT (U/L) | 27.5 (20- 41) |  |  |
| Albumin (g/dl) | 3.0 (2.9 - 3.4) |  |  |

\*: Statistically significant at $p \leq 0.05$ , BMI: body mass index, OSA: obstructive sleep
apnea, IHD: ischemic heart disease, PYI: pack/year index, PaO<sub>2</sub>: arterial partial
pressure of oxygen, PaCO<sub>2</sub>: arterial partial pressure of carbon dioxide, HCO<sub>3</sub>:
bicarbonate, SaO<sub>2</sub>: oxygen saturation, FBS: fasting blood sugar, Hb: hemoglobin,
BUN: blood urea nitrogen, Cr: creatinine, Na: sodium, K: potassium, AST: aspartate
transferase, ALT: alanine transferase.

### Diaphragmatic neural function assessment

Both CMEPs and PMEPs of studied population are illustrated in table “2” with demonstration example in figure “1”. There was statistically significant bilateral decrease in DRMT, CMEPA and PMEPA in COPD group versus healthy subjects (*p* < 0.0001 for all and 0.001 for PMEPA on right). Further, there was statistically significant increase in CMEPL and PMEPL bilaterally in COPD group versus healthy subjects (*p* < 0.0001 for all and 0.006 for CMEPL on right side). Left CMCT was significantly prolonged in COPD group vs. healthy subjects (*p* < 0.0001) but not for right CMCT “*p*= 0.376; table 2”. Further, there was significant increase in CMEPL and CMCT of left versus right diaphragm in COPD group “*p*= 0.003 and 0.001 respectively; table 3” but there was no statistically significant difference in control group.

**Figure 1.**
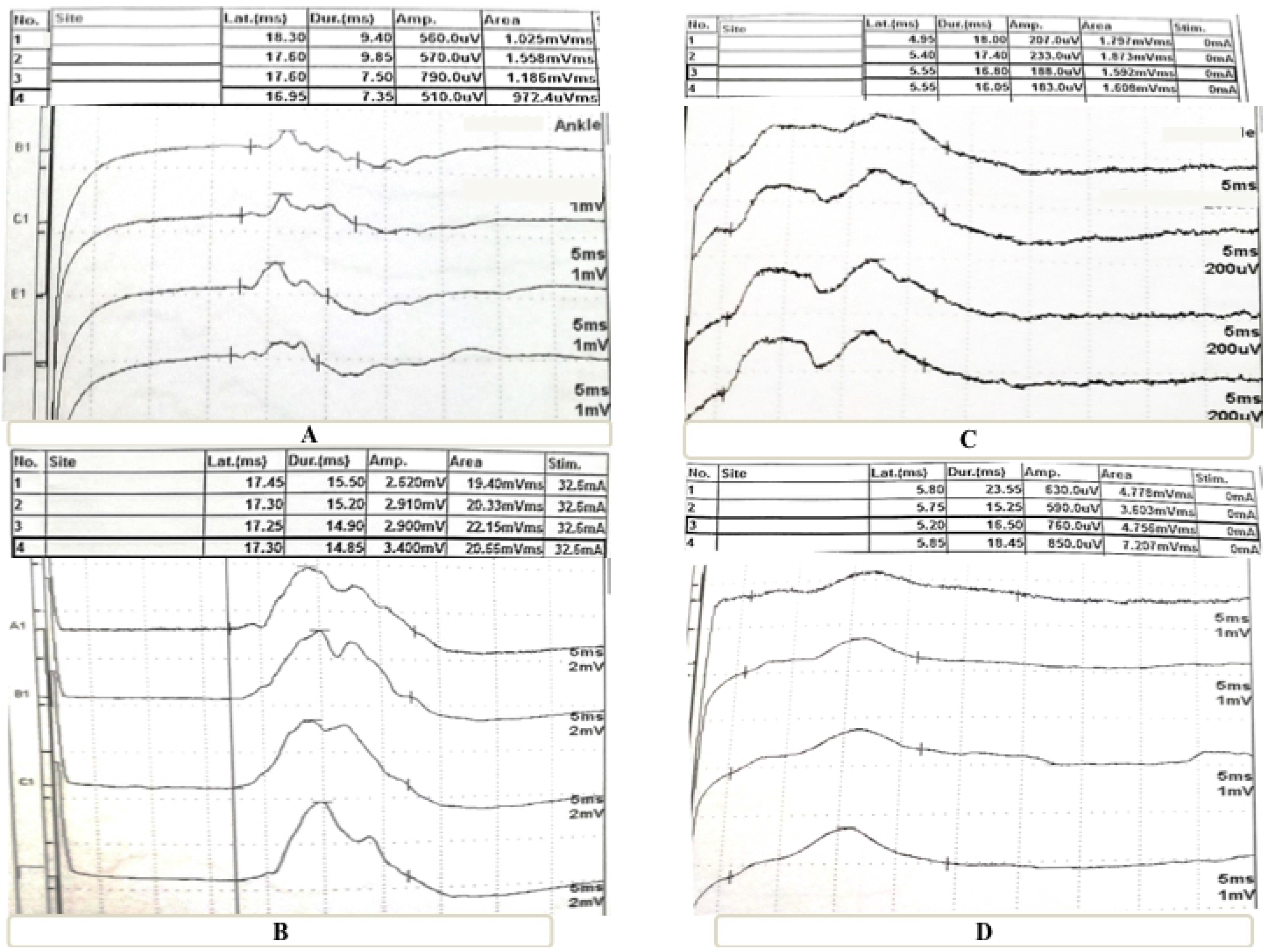
A: CMEP of diaphragm in COPD patient noticing that there is delayed latency and low amplitude of the response versus figure 1-B which represents healthy subject; C: PMEP of diaphragm in COPD patient with low amplitude of the response versus figure 1-D which represents healthy subject.

**Table (2):** Comparison between the two studied groups regarding diaphragmatic CMEP and PMEP parameters

| Parameter | COPD (n=30) | Control (n=17) | *p* value |
| --- | --- | --- | --- |
| **Right diaphragm conduction** |  |  |  |
| DRMT (%) | 66.9 ± 8.2 | 89.5 ± 5.2 | < 0.0001 |
| CMEPL (ms) | 14.4 (11.9 - 16.5) | 11.2 (10.5 - 12.4) | 0.006 |
| CMEPA (μv) | 120 (110-140) | 177 (158.3 - 180.0) | < 0.0001 |
| PMEPL (ms) | 6.99 ± 1.05 | 5.4 ± 0.6 | < 0.0001 |
| PMEPA (μv) | 135.0 (117.0 - 160.0) | 190 (179.5 - 196.3) | < 0.0001 |
| CMCT (ms) | 7.7 (4.9 - 9.2) | 5.9 (5.6 - 6.6) | 0.376 |
| **Left diaphragm conduction** |  |  |  |
| DRMT (%) | 68.6 ± 7.6 | 89 ± 4.4 | < 0.0001 |
| CMEPL (ms) | 16.8 (14.5 - 18.0) | 10.9 (10.6 - 12.8) | < 0.0001 |
| CMEPA (μv) | 127.1 ± 23.8 | 173.9 ± 34.2 | < 0.0001 |
| PMEPL (ms) | 7.4 (6.0 - 8.4) | 5.1 (4.7 - 5.75) | < 0.0001 |
| PMEPA (μv) | 147.3 ± 21.7 | 183.0 ± 35.9 | 0.001 |
| CMCT (ms) | 9.3 (8.1 - 10.1) | 6.2 (5.5 - 6.95) | < 0.0001 |

\*: Statistically significant at $p \leq 0.05$ , DRMT: diaphragmatic resting motor threshold, CMEPL: cortical motor evoked potential latency in milliseconds (ms), CMEPA: cortical motor evoked potential amplitude in microvoltage (μv), PMEPL: peripheral motor evoked potential latency, PMEPA: peripheral motor evoked
potential amplitude, CMCT: central motor conduction time.

**Table (3):** Comparison between right and left diaphragmatic CMEP and PMEP in both groups

| Parameter | COPD group (n=30) |  |  | Control group (n=17) |  |  |
| --- | --- | --- | --- | --- | --- | --- |
|  | Right | Left | *p* value | Right | Left | *p* value |
| **DRMT (%)** | 66.9 ± 8.2 | 68.6 ± 7.6 | 0.417 | 89.5 ± 5.2 | 89 ± 4.4 | 0.778 |
| **CMEPL (ms)** | 14.4 (11.9 - 16.5) | 16.8 (14.5 - 18.0) | 0.003* | 11.2 (10.5 - 12.4) | 10.9 (10.6 - 12.8) | 0.783 |
| **CMEPA (μv)** | 122.8 ± 22.3 | 127.1 ± 23.8 | 0.472 | 177 (158.3 - 180.0) | 173.9 ± 34.2 | 0.959 |
| **PMEPL (ms)** | 6.99 ± 1.05 | 7.4 (6.0 - 8.4) | 0.427 | 5.2 (4.9 - 5.8) | 5.1 (4.7 - 5.8) | 0.593 |
| **PMEPA (μv)** | 138.3 ± 25.7 | 147.3 ± 21.7 | 0.147 | 190 (179.5 - 196.3) | 190 (147.5 - 196.5) | 0.986 |
| **CMCT (ms)** | 7.7 (4.9 - 9.2) | 9.3 (8.1 - 10.1) | 0.001* | 5.9 (5.6 - 6.6) | 6.2 (5.5 - 6.95) | 0.629 |

\*: Statistically significant at $p \leq 0.05$ , DRMT: diaphragmatic resting motor threshold, CMEPL: cortical motor evoked potential latency in milliseconds (ms), CMEPA: cortical motor evoked potential amplitude in microvoltage (μv), PMEPL: peripheral motor evoked potential latency, PMEPA: peripheral motor evoked potential amplitude, CMCT: central motor conduction time.

## Correlations

Left diaphragmatic CMEPL and CMCT inversely correlated with different pulmonary function parameters (i.e. FVC% predicted, FEV_1_% predicted and FEV_1_/FVC) and positively correlated with CMEPA among the studied population “*p* < 0.01; figures 2A-F”. However, right diaphragmatic CMCT did not correlate with pulmonary function parameters (*p* > 0.05) while right CMEPL is inversely correlated with FVC% predicted (*p*= 0.036) but not FEV_1_% predicted or FEV_1_/FVC (*p* > 0.05) among the studied population. Both right and left diaphragmatic peripheral conduction (PMEPL and PMEPA) were positively correlated with different pulmonary function parameters (*p*< 0.01).

**Figure 2.**
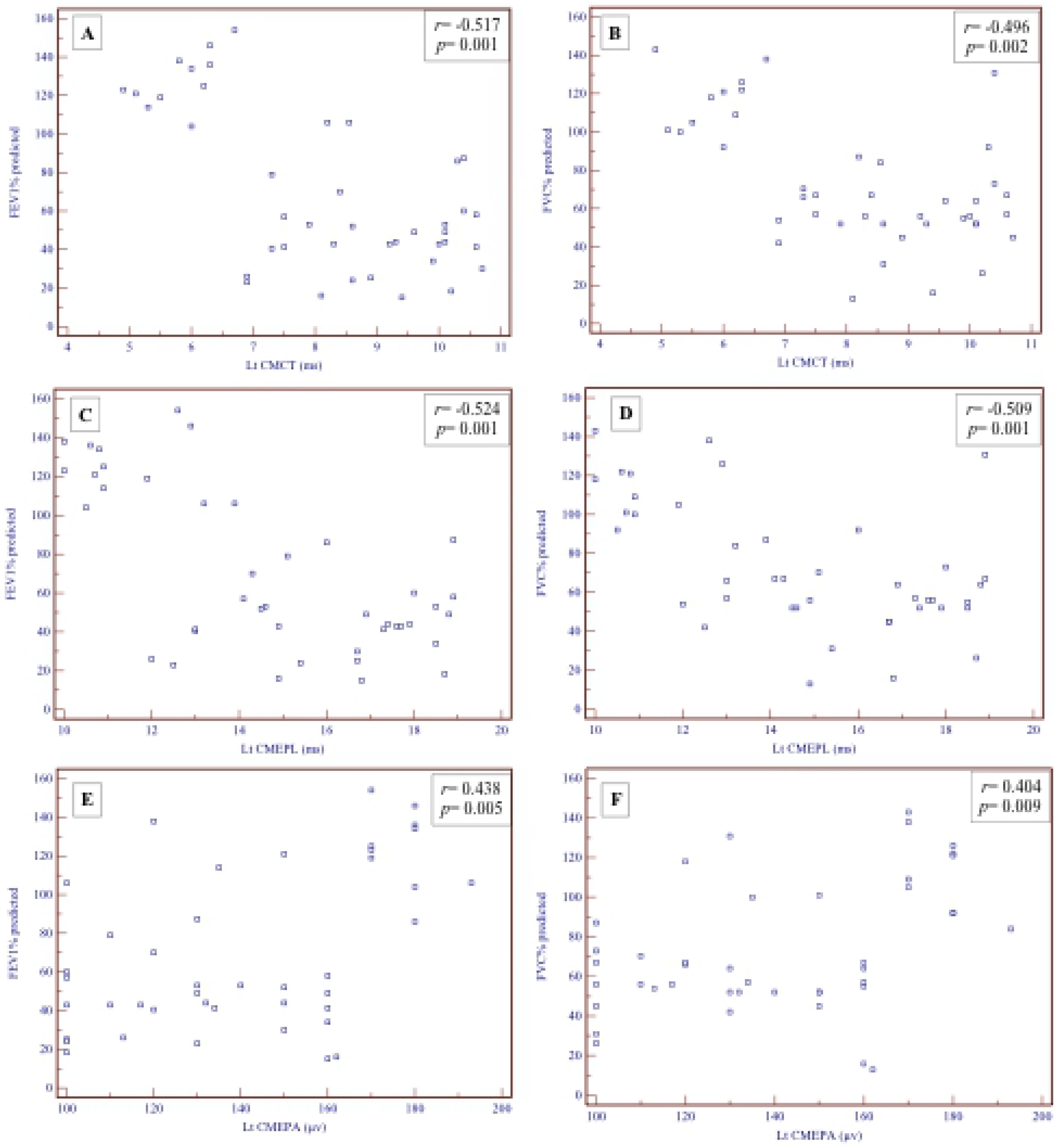
Correlations between spirometric parameters (FEV% predicted and FVC% predicted) and left CMEPs (A-F).

On the other hand, there was no statistically significant association between either CMEPs or PMEPs and COPD severity according to GOLD classification (*p* > 0.05). Similarly, there was no statistically significant correlation between diaphragmatic CMEPs or PMEPs and age, smoking status, smoking index, BMI, serum albumin or ABG parameters (*p* > 0.05).

### ROC analysis

According to ROC analysis, DRMT ≤80% had diagnostic accuracy of 98.6% to differentiate COPD from healthy control individuals with a sensitivity of 92% and specificity of 94% “AUC= 0.986, CI95%= 0.936 – 0.998, *p*= 0.0001; figure 3A”; CMEPL > 12.9 ms had diagnostic accuracy of 83% and sensitivity of 77% and specificity of 85% for differentiating COPD from healthy subjects “AUC= 0.828, CI95%= 0.737 – 0.898, *p*= 0.0001; figure 3B”; CMCT > 6.7 ms had diagnostic accuracy of 71.5% and sensitivity of 77% and specificity of 80% for differentiating COPD from healthy subjects “AUC= 0.715, CI95%= 0.612 – 0.803, *p*= 0.0001; figure 3C”; CMEPA ≤ 160 μv had 92% diagnostic accuracy, 98% sensitivity and 73.5% specificity for differentiating COPD from healthy subjects “AUC= 0.916, CI95%= 0.841 – 0.963, *p*= 0.0001; figure 3D”.

**Figure 3.**
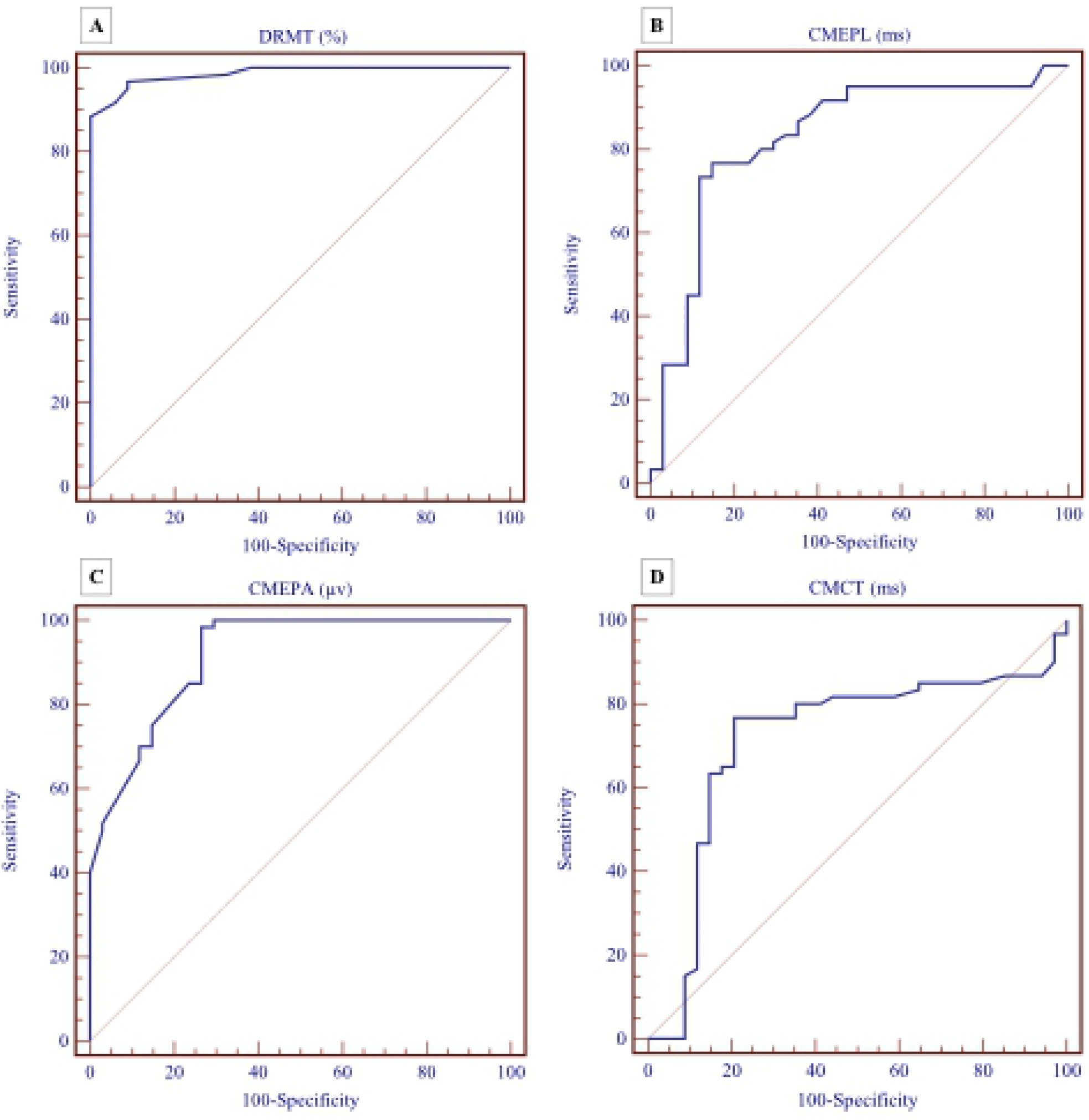
ROC analysis in COPD patients for predicting cutoff for CMEPs; A: for DMRT% (AUC= 0.986, CI95%= 0.936 – 0.998, *p*= 0.0001); B: for CMEPL (AUC= 0.828, CI95%= 0.737 – 0.898, *p*= 0.0001); C: CMEPA (AUC= 0.715, CI95%= 0.612 – 0.803, *p*= 0.0001); D: CMCT (AUC= 0.916, CI95%= 0.841 – 0.963, *p*= 0.0001).

## Discussion

In the current study, COPD patients had significant delayed central and peripheral diaphragmatic conduction latencies compared to the healthy control group, as well as decreased amplitude that was correlated with several parameters of pulmonary function testing. In addition, there was a statistically significant difference in COPD patients between right and left central diaphragmatic conduction.

### Previous studies and interpretation of the results

Hopkinson et al [18] found that diaphragmatic cortical motor thresholds were significantly lower in COPD than healthy controls as well as significant longer mean PMEPL. Similarly, Hamed et al [21] reported bilateral increase in CMEPL and CMCT in their studied COPD compared to healthy control group as well as decreased DRMT. Further, El-Tantawi et al [22] found peripheral phrenic nerve conduction abnormalities in 42.5% of their studied COPD patients that did not correlate with disease severity. These results are in accordance of the current results and could be explained by increased excitation of motor cortex and corticospinal pathways to the respiratory muscles in the COPD patient [23] and less excitability of intracortical facilitatory circuits at long interstimulus intervals using paired stimulation denoting ceiling effect of motor control output to the respiratory muscles of case of COPD. [18]

Interestingly, we found significant increase in CMEPL and CMCT of left versus right diaphragm in COPD group which correlated inversely with FEV_1_% and FVC% but not ABG parameters. This denotes that there is heterogeneity in affection of respiratory muscles which is in accordance with disease heterogeneity on one hand. [24] On the other hand, increased inspiratory load of respiratory muscles has been associated with significant activation of several motor cortical areas as demonstrated by increased regional cerebral blood flow using positron emission tomography [25] which could be affected asymmetrically. More recently, Dodd et al [26] demonstrated by magnetic resonance imaging techniques that generalized functional activation of resting-state networks in COPD patients compared with controls.

Further, we proposed cutoff point for CMEPs that had good diagnostic accuracy and sensitivity for predicting corticospinal pathway affection in case of COPD. Lissens [8] demonstrated values for diaphragmatic CMEPs in 10 healthy man only. However, to our knowledge, there are no specific values proposed to date that could be reference for CMEPs responses in COPD. We suppose that the presented values could be considered as reference, however, further studies with larger population should be considered to confirm the current values.

### Clinical implementation

Diaphragmatic dysfunction is strongly correlated with FEV_1_ in COPD [27] and correlated with the perception of dyspnea among this group of patients. [28] Coticospinal pathway affection could be another factor for development of diaphragmatic dysfunction in COPD patients accordingly its evaluation could help in personalization of COPD management especially pulmonary rehabilitation programs. Chun et al found significant improvement of diaphragmatic motility after pulmonary rehabilitation using sonography. [29]

Further, assessment of diaphragmatic corticospinal pathway could be of value in evaluation noninvasive ventilation use in stable severe/ very severe COPD. [30] This has been demonstrated by Hopkinson et al, [23] who found that the excitability of the corticospinal pathway to the diaphragm reduced in 6 COPD patients after acute noninvasive ventilation use. This could be explained by the fact that noninvasive ventilation reduced inspiratory muscles loads [30] through reduces the cortical motor areas excitability supplying the respiratory muscles especially the diaphragm. [31] Accordingly, TMS could be a good applicable tool for evaluation of central and peripheral diaphragmatic neural pathway which may affect the management of COPD patients.

### Limitations

The current study has some limitations. Firstly, we studied only the diaphragm as the main respiratory muscle and we did not study the intercostals or abdominal muscles. Further, the cortical area for the diaphragm has been previously validated in healthy man [9, 12] rather than other respiratory muscles. Secondly, we used surface electrodes for diaphragm CMEPs recording and we did not use diaphragm needle electromyography. However, intercostal surface electrodes have been validated previously [19] and needle electromyography is more invasive and could be associated with complications as pneumothorax. Lastly, we did not study the diaphragmatic CMEPs response at different intervals of time of at maximal inspiratory efforts in COPD patients. However, Sharshar et al [32] studied before the response to cortical stimulations at different points of time or inspiratory efforts in healthy men and they concluded that cortical motor control of diaphragm is identical during different inspiratory tasks.

## Conclusions

Central cortico-diaphragmatic motor system is affected in COPD patients with heterogeneity of both sides that is correlated with airway obstruction as being detected with spirometry but not with COPD severity or ABG changes. The cutoff values for CMEPs in COPD patients in the current study had good diagnostic value in predicting diaphragmatic corticospinal affection. The current data could be a step for future studies for evaluating the diaphragm using noninvasive tool - the TMS - after therapeutic interventions for COPD.

## Acknowledgments

The authors would like to thank the technicians/nurses in the pulmonary function unit for their willing to help in the current manuscript. Dr. Shafiek is the guarantor of the manuscript.

